# Catching thoughts: self-caught experience sampling preferentially captures characteristic features of off-task experiences across the life span

**DOI:** 10.1101/2022.12.15.520676

**Authors:** Léa M. Martinon, Jonathan Smallwood, Leigh M. Riby

## Abstract

Understanding transient states, like off-task mind-wandering, is assumed to be improved by capitalizing on our ability to recognize changes in our stream of thought, a process known as meta-awareness. We test this assumption by comparing mind-wandering content when noticed by the participant (self-caught) against those thoughts reported after externally initiated probes (probe-caught). Thirty-eight older and 36 younger individuals completed a cognitive task. At the same time, multiple feature descriptions of thoughts (task-relevance, temporal focus, and self-referential) were captured using self and probe-caught methods. Using a pattern-learning approach, we established that self-caught experiences produce similar but generally “noisier” estimates compared to those reported at probes. However, self-caught experiences contained more off-task characteristics relative to reports at probes. Importantly, despite reductions in off-task thought, older adults retain the ability to self-catch experiences with these features. Our study establishes self-catching ability as an essential means of revealing the detailed content of off-task states, an ability relatively well maintained into old age.

## Introduction

Patterns of ongoing thoughts are heterogeneous in their content, varying over time, across contexts and people (Smallwood et al., 2021). Off-task mind-wandering entails individuals’ attention becoming momentarily decoupled from sensory input and focused on self-generated thoughts and feelings (e.g. Barron et al. 2011). It has been argued that a clearer understanding of such transient conscious states can be achieved by considering how we take stock of changes that occur in our stream of thought, a process known as meta-awareness (Allen et al., 2013; Schooler, 2002). One core assumption is that introspection allows individuals to notice spontaneous changes in their conscious experiences, which can be experimentally captured by comparing experiences that individuals notice (self-caught) compared to when experiences are reported at experimenter-initiated periods (probe-caught; Smallwood & Schooler, 2006). Prior studies have combined the probe and self-caught methods to show that alcohol intoxication (Sayette et al., 2009) and cigarette craving (Sayette et al., 2010) tend to increase the probability of mind-wandering experiences (probe-caught) without necessarily changing the probability of noticing them (self-caught). Studies of this nature provide evidence that probe-caught sampling captures experiences individuals’ have at least temporarily failed to recognize. In contrast, self-caught experiences result from the capacity to notice specific features of cognition (which may be compromised when drunk through alcohol). Our study seeks additional evidence for the meta-awareness theory by examining whether self-catching provides insight into experiential features characteristic of spontaneous states in a way that is different from reports provided at probes. This endeavour would be consistent with the suggestion from early work in the area arguing for distinct, separable processes involved in the generation of mind-wandering episodes and the awareness of such events (e.g. Sayette et al., 2009).

Patterns of ongoing experience vary substantially with age. Older individuals report patterns of task-focused thoughts more commonly than younger individuals (Jordão et al., 2019; Maillet & Schacter, 2016). A meta-analysis concurs with less frequent mind-wandering episodes reported for older compared to younger adults (Hedges’ g=-.89; Jordão et al., 2019), and this effect is even more pronounced as we enter late adulthood. However, the specific factors which underlie age-related changes in thought patterns remain a matter of debate. One possibility is that structural brain changes (e.g. Damoiseaux, 2017; Fjell et al., 2009; West, 2000) in older individuals lead to insufficient cognitive resources to perform a task. Consequently, they must prioritize resources to maintain task-relevant information processing. However, one view, even in the face of these difficulties, is that older individuals can compensate for age-related changes through increased reliance on crystallized rather than fluid intelligence (Spreng et al., 2018). It has also been suggested that age-related differences in motivation mediate this relationship (Frank et al., 2015; Krawietz et al., 2012; Seli et al., 2020; Shake et al., 2016). For example, older adults are more concerned about their performance than young adults (e.g. stereotype threat; Jordano & Touron, 2017), which can affect their task engagement (Ennis et al., 2013). Finally, a lack of familiarity with testing typically occurring in university contexts may explain why older adults engage in more on-task experiences (McVay et al., 2013; McVay & Kane, 2010). Despite the different reasons why ongoing thought patterns change with age, our study compared older and younger individuals to examine whether the capacity to notice off-task episodes in experience is preserved in a population from whom off-task states are reduced.

Our current study, therefore, had two aims. First, we tested whether the capacity for meta-awareness, defined by unique experiential features derived from self-initiated experience sampling, provides important insight into transient changes in experience (off-task mind-wandering). Second, we examined whether the ability to take stock of specific experience features is retained into old age when we know that off-task states are reduced. To capture different thought patterns, we used multi-dimensional experience sampling (MDES; Smallwood et al., 2016). In this method, participants are asked to describe their experience on several dimensions, and then data are decomposed into the latent patterns that underly the reports. This method previously described patterns that capture detailed, evolving, deliberate task-relevant thoughts, off-task social episodic thoughts, and verbal and self-relevant thoughts. These different thought patterns have been linked to unique neural substrates (Konu, Turnbull, et al., 2020; Sormaz et al., 2018; Turnbull et al., 2019).

Combining MDES with self- and probe-caught measures of experience allowed us to employ pattern learning methods to assess the possibility that self-caught experiences contain more information regarding transient changes in experience than reports of experiences provided by probes (Schooler et al., 2011). To achieve this goal, we analyzed our data in two different ways. First, we examined qualitative differences between patterns of thought captured by self- and probe-caught experience sampling. Here, we derived patterns of experience from both self- and probe-caught methods separately and compared these to those seen across the entire data set. This allows us to establish the degree to which patterns formed by each method yield descriptions of experience which differ from those seen when analyzed collectively. Second, we examined whether the different methods of experience sampling capture ‘quantitative’ differences in a common set of experiences. In this analysis, we derived common patterns of experience using data from both methods and examined whether moments characterized by different methods differentially contribute to this common ‘experiential’ space (Ho et al., 2020).

## Method

### Participants

A convenience sample of 36 young adults (*M* = 21.94, *SD* = 4.49; women = 29) and 38 older adults (*M* = 69.42, *SD* = 7.42; women = 24) were recruited for this study. The sample size was determined based on similar studies of this nature and anticipated large effect sizes (see Martinon et al., 2019). The young and older adults’ mean years of education were 15.71 (*SD* = 2.63) and 16.20 (*SD* = 4.13), respectively. Participants received either course credits or gift vouchers in compensation for their time and travel. Inclusion criteria were to be a native English speaker and to have normal or corrected vision and hearing. Exclusion criteria were the presence or history of a neurological or psychiatric disorder, currently taking antidepressants, and regular practice of meditation, Tai-chi, or Yoga in the past 3 years. Older participants completed the Mini-Mental State Examination (MMSE, Folstein et al., 1975) to ensure that they did not have dementia or mild cognitive impairment (threshold: score ≥ 26/30; *M* = 28.63, *SD* = 1.28). This study was approved by the Ethics Committee of the Faculty of Health and Life Sciences of Northumbria University. The investigation was conducted according to the principles expressed in the Declaration of Helsinki and participants provided written informed consent.

### Vigilance task

The vigilance task was developed using PsychoPy (Peirce, 2007) and featured a 0-Back procedure based on the task developed by Konishi et al. (2015). Participants were presented with different pairs of shapes (non-targets) appearing on the screen divided by a vertical line; the pairs could be: a circle and a square, a circle, and a triangle, or a square and a triangle, with a total of 6 possible pairs (two different left/right configurations for each). The pairs never had shapes of the same kind (e.g. a square and a square). A block of non-targets was followed by a target requiring participants to make a manual response. The target was a small stimulus (either a circle, square, or triangle) presented in the centre of the line, in blue. Two shapes flanked the target, and participants had to indicate, by pressing the left or right arrow key, on which side was the same shape as the target shape (see the upper panel of Figure 1). Every presentation of non-targets, targets and probe screens was separated by a fixation cross. The fixation crosses, non-targets, and targets were respectively presented for 1.5, 1, and 2 seconds and a response from participants did not end the target presentation. Targets were presented randomly with a ratio of 1 target for 5 non-targets. The task was designed to last 15 minutes (with the exclusion of the time taken to answer probes) which resulted in an average of 60.95 (SD=7.47) targets.

**Figure 1.**
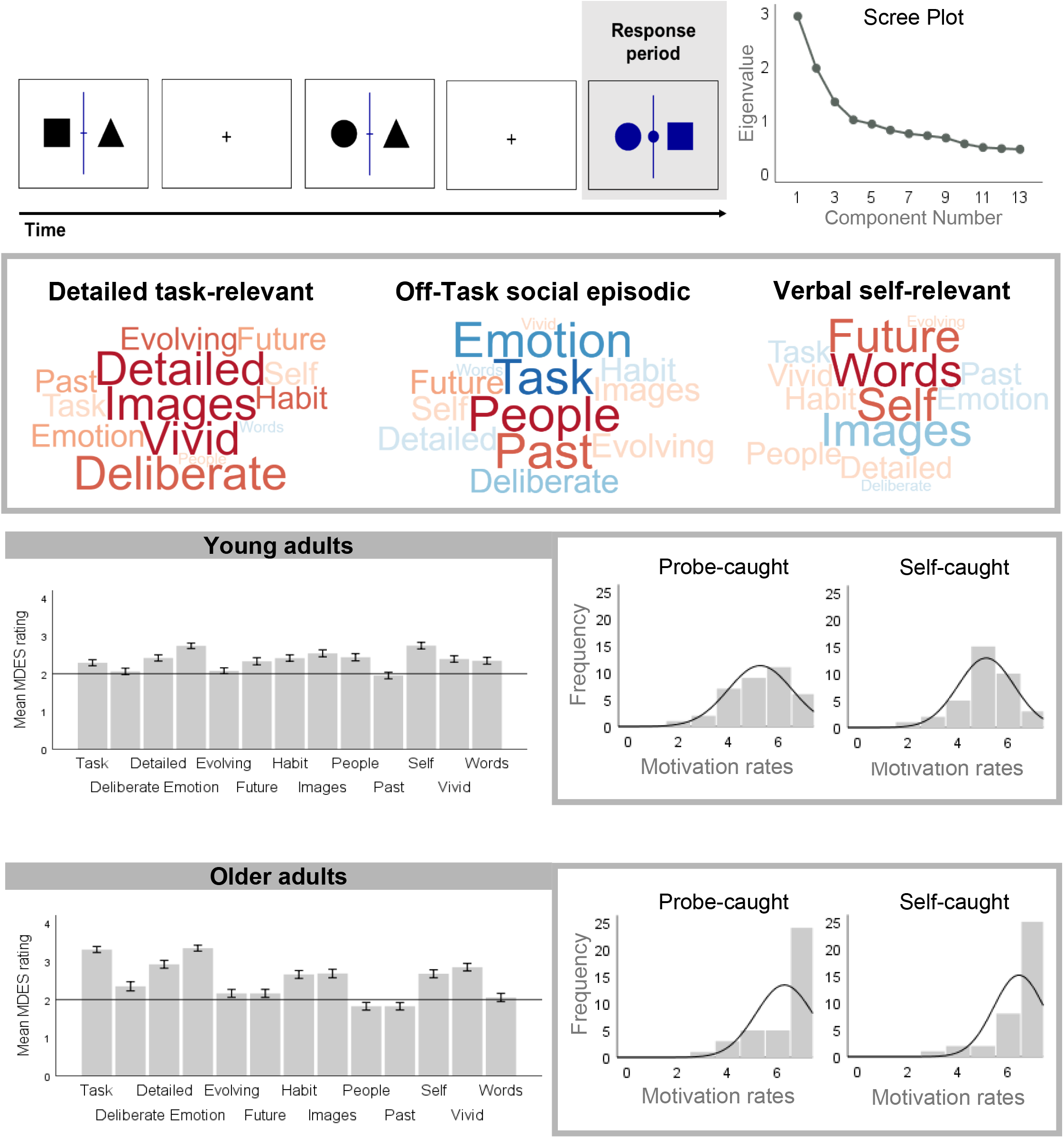
In the left upper panel is an illustration of the 0-Back task used. Note that the correct response of the example target trial is “circle”. Decomposition of the experience sampling data collected during this task revealed three components across all conditions. Based on their loadings the three components were summarised as “Detailed task-relevant”, “Off-Task social episodic”, and “Off-task verbal self-relevant”. The word clouds summarise these loadings in which the colour of the word describes the direction of the relationship (red = positive, blue = negative) and the size of the item reflects the magnitude of the loading. The scree plot for this decomposition is presented on the right upper panel. Bar-plots in the left lower panel show the mean ratings for each item that these components are derived from, for both young and older adults. Error bars represent 99.5% Confidence Intervals, bonferoni corrected for the number of questions. Histograms in the right lower panel represent the distribution of motivation rates for younger and older adults, for both probe- and self-caught measures.

The Multi-Dimensional Experience Sampling method was employed to monitor the participant’s experience while completing the abovementioned task (Smallwood et al., 2016). Participants completed two blocks - one with a probe-caught method and one with a self-caught method, using a counterbalanced design. In both cases, thought probes consisted of an on-screen prompt for the participants to rate their focus level (‘*My thoughts were focused on the task I was performing*.’) on a four-point Likert scale from 1 (‘*Not at all*’) to 4 (‘*Completely*’), using numbers on the keyboard. This prompt was followed by 12 questions regarding the characteristics of their thoughts immediately before the probe (Table 1.). Probes were presented randomly with a ratio of 1 probe for 13 trials with a maximum of 20 probes. An average of 19.39 (*SD* = 1.60) probes was presented. In the self-caught method, participants were asked to carefully monitor their thoughts during the task and press the ‘p’ key every time they noticed their mind had drifted away. When pressing the ‘p’ key the same 13 questions were automatically presented on the screen. On average, participants pressed the ‘p’ key 8.91 times (*SD* = 8.12). In both conditions, the task was set to last 15 minutes, excluding the time taken to answer the questions. The full completion of each task varied between 20 and 30 minutes.

**Table 1.** Questions and scales used in Multi-dimensional Experience Sampling.

| Measures | Probe questions | Scale (0 → 4) |
| --- | --- | --- |
| Task | My thoughts were focused on the task I was performing. | Not at all → Completely |
| Future | My thoughts involved future events. | Not at all → Completely |
| Past | My thoughts involved past events. | Not at all → Completely |
| Self | My thoughts involved myself. | Not at all → Completely |
| People | My thoughts involved other people. | Not at all → Completely |
| Emotion | The content of my thoughts was: | Negative → Positive |
| Images | My thoughts were in the form of images. | Not at all → Completely |
| Words | My thoughts were in the form of words. | Not at all → Completely |
| Vivid | My thoughts were vivid as if I was there | Not at all → Completely |
| Detailed | My thoughts were detailed and specific. | Not at all → Completely |
| Habit | This thought has recurrent themes similar to those I have had before. | Not at all → Completely |
| Evolving | My thoughts tended to evolve in a series of steps | Not at all → Completely |
| Deliberate | My thoughts were: | Spontaneous → Deliberate |

### Procedure

Participants were seen between one hour and thirty minutes and two hours in the laboratory. At the beginning of the session, all participants gave informed consent, followed by a set of demographic questions. Older adults also completed the MMSE. Participants were then asked to complete the vigilance task. The task was completed once with a probe-caught sampling method and once with a self-caught sampling method. The order of the presentation of the tasks was randomized. Before each task participants were required to rate their motivation to perform the task using a Likert scale from 1 (Not at all motivated) to 7 (Extremely motivated). All participants completed a practice trial of the task beforehand.

## Results

### Behavioural performance

Differences in participants’ performance on the task were investigated according to age (young vs. older) and type of measure (probe-caught vs. self-caught). First, a repeated-measure Analysis of Variance (ANOVA) was conducted on participants’ accuracy scores, showing a main effect of measure with anecdotal Bayesian evidence in favour of this effect [*F*(1, 71)=5.553, *p*<.05, *η*_*p*_*²*=0.07, BF_10_^1^=1.779, BF_01_=0.562], illustrating worse performance in the self-caught method compared to the probe-caught method (see Table 2.). This makes sense as the self-caught method requires additional cognitive resources for participants to monitor their thoughts and notify any mind drifts. This analysis suggests that these supplementary demands can impact performance on the ongoing task. However, there were no effect of age [*F*(1, 71)=.058, *p*=.810, *η*_*p*_*²*=0.00, BF_10_=0.330], nor an interaction effect between age and type of measure with anecdotal Bayesian evidence in favour of this effect [*F*(1, 71)=3.42, *p*=.069, *η*_*p*_*²*=0.05, BF_10_=1.073^2^, BF_01_=0.933]. See Table S1 in the supplementary materials for a comparison of the models in a Bayesian repeated-measure ANOVA.

**Table 2.**
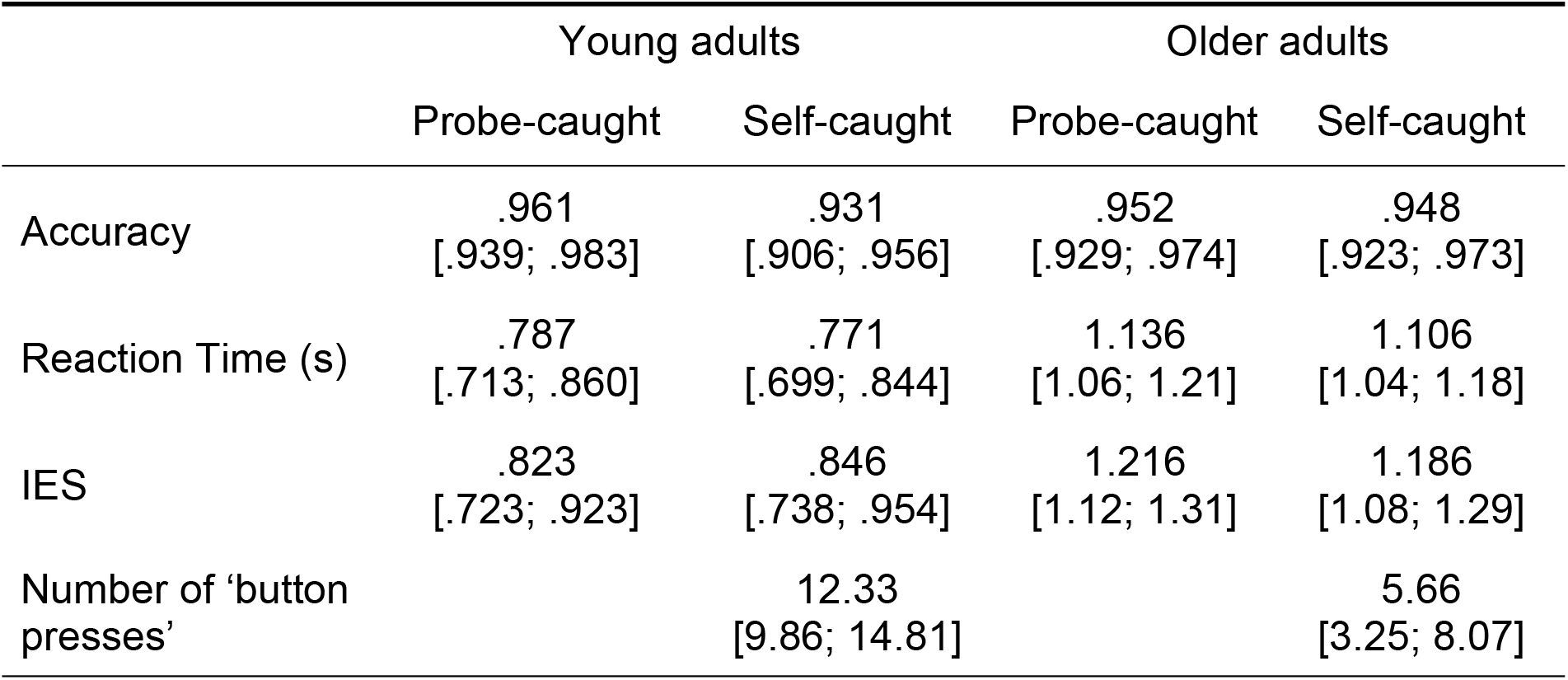
Descriptive statistics, means [95% confidence intervals], of young and older adults on the vigilance task.

|  | Young adults |  | Older adults |  |
| --- | --- | --- | --- | --- |
|  | Probe-caught | Self-caught | Probe-caught | Self-caught |
| Accuracy | .961<br>[.939; .983] | .931<br>[.906; .956] | .952<br>[.929; .974] | .948<br>[.923; .973] |
| Reaction Time (s) | .787<br>[.713; .860] | .771<br>[.699; .844] | 1.136<br>[1.06; 1.21] | 1.106<br>[1.04; 1.18] |
| IES | .823<br>[.723; .923] | .846<br>[.738; .954] | 1.216<br>[1.12; 1.31] | 1.186<br>[1.08; 1.29] |
| Number of 'button presses' |  | 12.33<br>[9.86; 14.81] |  | 5.66<br>[3.25; 8.07] |

Next, we considered participants’ performance while accounting for processing speed. An inverse efficiency score (IES), representing the average energy consumed by a system over time, was computed following Townsend & Ashby (1983) recommendations: reaction time (RT) divided by the proportion of correct answers (PC): IES = RT/PC. Results of the repeated-measure ANOVA showed a main effect of age with extreme Bayesian evidence in favour of this effect [*F*(1, 71)=27.50, *p*<.001, *η*_*p*_*²*=0.28, BF_10_=6171.07], where younger adults performed the task more efficiently than older adults. There were no effect of type of measure [*F*(1, 71)=.026, *p*=.873, *η*_*p*_*²*=0.00, BF_10_=0.182], nor interaction effect between age and type of measure [*F*(1, 71)=1.48, *p*=.227, *η*_*p*_*²*=0.02, BF_10_=0.422] (see Table 2 for descriptive data for these comparisons). See Table S2 in the supplementary materials for a comparison of the models in a Bayesian repeated-measure ANOVA

Lastly, we investigated the effect of age on the number of times participants press the button during the self-caught blocks. Evidence from a one-way ANOVA shows that younger adults pressed the ‘p’ button more often than older adults, *F*(1, 72)=14.86, *p*<.001, *η*_*p*_*²*=0.17, BF_10_=100.24. This replicates prior reductions in off-task thoughts in older adults using the self-caught method (for a review see Jordão et al., 2019; and Maillet & Schacter, 2016). See Table S3 in the supplementary materials for a comparison of the models in a Bayesian ANOVA.

### Principal Component Analysis identifies three types of thoughts

Several participants did not report any mind-wandering experiences with the self-caught method. Thus, data from one young and 10 older adults have been excluded. The following analyses, therefore, will be conducted on data from 35 young adults and 28 older adults.

To examine the MDES data, the set of questions was decomposed using principal component analysis (PCA), applying varimax rotation (e.g. Ho et al., 2020; Karapanagiotidis et al., 2017; Konu, Mckeown, et al., 2020; Konu, Turnbull, et al., 2020; Martinon et al., 2019; Smallwood et al., 2016). This allowed the core patterns of variance within the self-reported data to be characterized in a smaller set of underlying dimensions. Three-factor solutions were selected with eigenvalues > 1, which are presented in the form of word clouds in Figure 1 (see Figure 1 for scree plot and Table S4. in supplementary materials for detailed factor loadings). <u>Component One</u>: *Detailed task-relevant thoughts* - with high weighting on items indicating detailed, deliberate and vivid thoughts accounted for 22.59% of the variance. <u>Component Two</u>: *Off-task social episodic thoughts* - with high weighting showing thoughts that tend to be about other people and the past while being negative and not about the task, accounted for 15.10% of the variance. <u>Component Three</u>: *Off-task verbal self-relevant thought* - with high weighting reflecting thoughts that were in the form of words (and not images) and about the self and the future, accounted for 10.26% of the overall variance.

### Qualitative differences in patterns of ongoing thoughts between groups and sampling methods

Having described the patterns that account for the most variance within the MDES data, we examined the qualitative similarity in thought patterns captured by different experience sampling methods and in different age groups. To achieve this goal, we generated new factor solutions while separating young and older adults’ data as well as the probe-caught and self-caught data (4 cells of our study). In these new PCAs we specifically asked for 3 components to maintain similarity with the original solution (see Ho et al., 2020 for a previous example of a similar analysis). Next, we compared these three new factors to the original omnibus solution described above, which used the combination of all the data regardless of age or method sampling (see Figure 1). This analysis allows us to evaluate the possibility of qualitative differences in the patterns of experience across methods or age groups by testing the null hypothesis that the PCA patterns captured by the omnibus analysis are only present in only some subsets of the data.

To evaluate the correspondence between the new factors and the omnibus factors, while accounting for the hierarchical structure of the data, linear mixed models were used. The MIXED procedure in SPSS 26.0 (Peugh & Enders, 2005) with Maximum Likelihood Estimation was used. For these analyses, all dependent and independent variables were grand mean centred to ensure clarity in interpreting any effects. All linear mixed models included one random effect (the intercept for each participant) to control for the dependency arising due to repeated sampling of data within participants. The omnibus factor was set as the outcome variable for each of the three components. Fixed predictors were the corresponding factor generated separately for each experimental condition (new factor: PCA loadings described above), participants’ age group (old or young), and the type of measure used (self or probe). We also included the following interactions: age group * new factor, measure * new factor, and age group * measure * new factor. To control family-wise errors resulting from the multiple comparisons for three components alpha levels were Bonferroni corrected to.017.

In these analyses, a main effect of new factor describes variance that is shared by the new PCAs and the omnibus solution (e.g. regardless of the method of report), the main effect of method describes the difference in the similarity between the omnibus PCA and the features seen when only focusing on one method, a main effect of age describes how similarity between the omnibus and the new PCA’s changes with age, and finally interactions with age group indicate that the similarity with PCA depends on the method of report used to collect experience sampling data. Supplementary Figure S1 shows the loadings for each of the PCAs calculated on each segment of the data in the form of word clouds, as well as the scree plots from each solution.

Analyses for **detailed task-relevant** thoughts showed that for every increase of one standard deviation of the new factor, the omnibus factor increases of .864 (*SE*=.008), independently of participants’ age or the method used, *t*(1483)=107.66, *p*<.001, CI98.3% [.845;.883]. The age group * new factor interaction establishes that, going from young to older adults, the effect of the new factor on the omnibus factor increased by .124 (*SE*=.016), regardless of method, *t*(1399)=7.66, *p*<.001, CI98.3% [.086; .163]. The measure * new factor interaction shows that, going from the probe-to the self-caught method, the effect of the new factor on the omnibus factor decrease of -.403 (*SE*=.016), regardless of age, *t*(2050)=-25.58, *p*<.001, CI98.3% [-.440; -.365].

Finally, the age group * measure * new factor interaction suggests that going from the probe-to the self-caught method, the difference in the relationship between the new factor and the omnibus factor between young and older adults is -.322 (*SE*=.032), *t*(2031)=-9.98, *p*<.001, CI98.3% [-.399; -.245] (see Figure 2.). To unpack this three-way interaction, we conducted linear mixed models separately for young and older adults. For young adults, going from the probe-to the self-caught method, led to a decrease in the similarity of the new factor with the omnibus factor of -.257 (*SE*=.020), *t*(1143)=-12.83, *p*<.001, CI98.3% [-.304; -.209]. In the probe-caught method, for every increase of one standard deviation of the new factor, the omnibus factor increased by .940 (*SE*=.006) while when using the self-caught method it increased by .620 (*SE*=.022). For older adults, going from the probe-to the self-caught method, the similarity between the new factor and the omnibus factor decreased by -.549 (*SE*=.020), *t*(856)=-27.45, *p*<.001, CI98.3% [-.597; -.501]. In the probe-caught method, for every increase of one standard deviation of the new factor, the omnibus factor increased by 1.13 (*SE*=.009), while in the self-caught method it increased by .601 (*SE*=.028). Overall, patterns of detailed task-relevant thought are present in both old and young individuals using both methods of report. However, better correspondence between this pattern was observed in the reports provided by older rather than younger adults, and when experiences were reported via probes rather than when self-caught.

**Figure 2.**
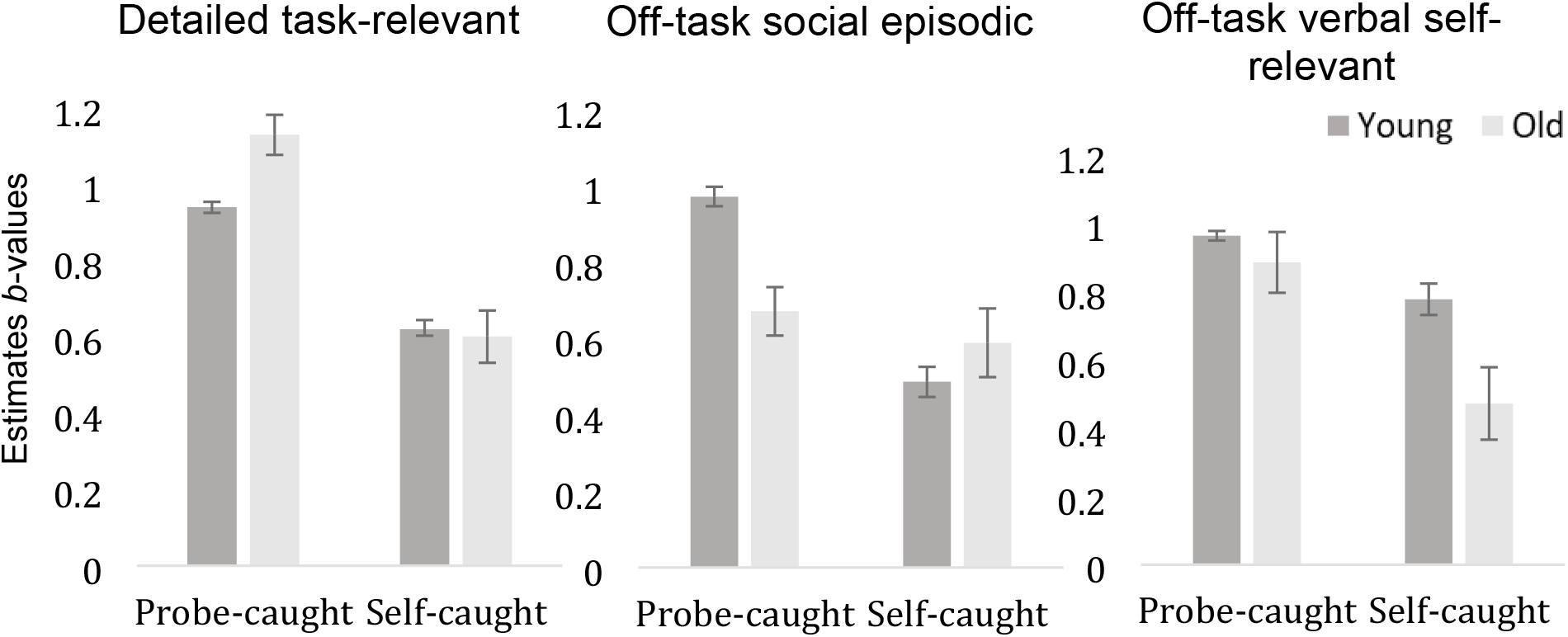
Estimate *b*-values illustrating the contribution of PCA components generated separately for age groups and type of measures onto the omnibus PCA components for each group and type of measure. Error bars represent 98,3% Confidence Intervals, bonferoni corrected for the number of factors.

Analyses for **off-task social episodic** thoughts showed that for every increase of one standard deviation of the new factor, the omnibus factor increased by .707 (*SE*=.010), independently of participants’ age or the method used, *t*(2094)=70.87, *p*<.001, CI98.3% [.683; .731]. The age group * new factor interaction indicated that going from young to older adults, and regardless of method, the similarity between the new factor and the omnibus factor decreased by -.174 (*SE*=.020), *t*(2094)=-8.70, *p*<.001, CI98.3% [-.222; -.126]. The measure * new factor interaction indicates that, regardless of age going from the probe-to the self-caught method, the similarity between the new factor and the omnibus factor decreased by -.312 (*SE*=.021), *t*(2087)=-14.76, *p*<.001, CI98.3% [-.362; -.261]. Finally, the age group * measure * new factor interaction indicates that going from the probe-to the self-caught method, the similarity between the new factor and the omnibus solution decreased from younger to older individuals by .295 (*SE*=.043), *t*(2088)=6.83, *p*<.001, CI98.3% [.192; .398] (see Figure 2.). We conducted linear mixed models separately for young and older adults, which established that for young adults, going from the probe- to the self-caught method reduced the similarity between the new factor and the omnibus factor by -.441 (*SE*=.026), *t*(1144)=-17.27, *p*<.001, CI98.3% [-.502; -.380]. In the probe-caught method, for every increase of one standard deviation of the new factor, the omnibus factor increased by .972 (*SE*=.011) while it increased by .487 (*SE*=.027) with the self-caught method. For older adults, going from the probe-to the self-caught method, the effect of the new factor on the omnibus factor decreased by -.153 (*SE*=.034), *t*(942)=-4.53, *p*<.001, CI98.3% [-.235; -.072]. In the probe-caught method, for every increase of one standard deviation of the new factor, the omnibus factor increased by .672 (*SE*=.017) while it increased by .589 (*SE*=.037) with the self-caught method. Overall, patterns of off-task social episodic thought are well captured by patterns seen in both age groups and sampling methods, with the highest correspondence for the probe-caught method for younger adults.

Analyses for **off-task verbal self-relevant** thoughts indicated that, regardless of participants’ age or the method used, for every increase of one standard deviation of the new factor, the omnibus factor increases by .79 (*SE*=.012), *t*(1985)=65.36, *p*<.001, CI98.3% [.762; .820]. The age group * new factor interaction indicated that, regardless of method, going from young to older adults, the similarity between the new factor and the omnibus factor decreased by -.231 (*SE*=.024), *t*(1960)=-9.45, *p*<.001, CI98.3% [-.289; -.172]. The measure * new factor interaction suggests that, regardless of age, going from the probe-to the self-caught method, the effect of the new factor on the omnibus factor decrease of -.256 (*SE*=.024), *t*(2093)=-10.49, *p*<.001, CI98.3% [-.314; -.197]. Finally, the age group * measure * new factor indicates that going from the probe-to the self-caught method, the difference between young and older on the effect of the new factor on the omnibus factor decrease of -.204 (*SE*=.050 *t*(2092)=-4.101, *p*<.001, CI 98.3% [-.323; -.085] (see Figure 2.). We next conducted linear mixed models separately for young and older adults identifying that for young adults, going from the probe-to the self-caught method, the similarity between the new factor and the omnibus factor decreased by -.165 (*SE*=.027), *t*(1139)=-6.24, *p*<.001, CI98.3% [-.229; -.102]. In the probe-caught method, for every increase of one standard deviation of the new factor, the omnibus factor increased by .963 (*SE*=.006) while it increased by .777 (*SE*=.037) with the self-caught method. For older adults, going from the probe-to the self-caught method, the similarity between the new factor and the omnibus factor decreased by -.370 (*SE*=.043), *t*(950)=-8.57, *p*<.001, CI98.3% [-.473; -.267]. In the probe-caught method, for every increase of one standard deviation of the new factor, the omnibus factor increased by .885 (*SE*=.019) while with the self-caught method it increased by .472 (*SE*=.044). Overall, off-task verbal self-relevant thoughts are well captured by both age groups and sampling methods, however, correspondence is weakest using the self-caught method for older adults.

### Interim Summary

In each of the three thought patterns seen in our omnibus PCA we found a substantial amount of shared variance in the patterns generated based on each subset of the data (Range of Beta Values =.472–1.13). However, we also found that the patterns reported using the probe-caught method were generally higher (Range of Beta Values =.672– 1.13) than for the self-caught method (Range of Beta Values =.472–.777). This pattern suggests that the self-caught and probe-caught methods describe patterns with broadly similar features, yet the former provides “noisier” estimates than the latter. Notably, however, there was no evidence that one age group systematically contributed to the overall patterns of experience: detailed task focus was more clearly seen in older individuals reports (*b*_Young_=.806, *b*_Old_=.928), while off-task states were generally more clearly seen in younger individuals (Off-task episodic *b*_Young_=.787, *b*_Old_=.613; Off-task verbal *b*_Young_=.896, *b*_Old_=.662). Together, these analyses suggest that although there is significant variation in how specific cells in our study contribute to the patterns seen in the omnibus solution, there is also reasonable similarity across methods that allows us to reject the null hypothesis that experience sampling using self and probe-caught methods (or in older or younger individuals) necessarily capture qualitatively different patterns of experience.

### Quantitative differences in patterns of ongoing thoughts between groups and sampling methods

Having established broad similarities in the patterns of experience reported in each cell of our experiment, our next analysis examined quantitative differences between the patterns seen in each condition of our study and the overall dimensions described in the omnibus analysis. We conducted a repeated-measure ANOVA in which the average loading on the omnibus PCA for each individual in each cell were the dependent variables. We included repeated measures on Measure (Probe-caught, Self-caught) and a between-subject factor: Age (Old vs. Young). To control family-wise errors resulting from the multiple comparisons for three components, alpha levels were Bonferroni corrected to .017. The results of these analyses are presented in Figure 3.

**Figure 3.**
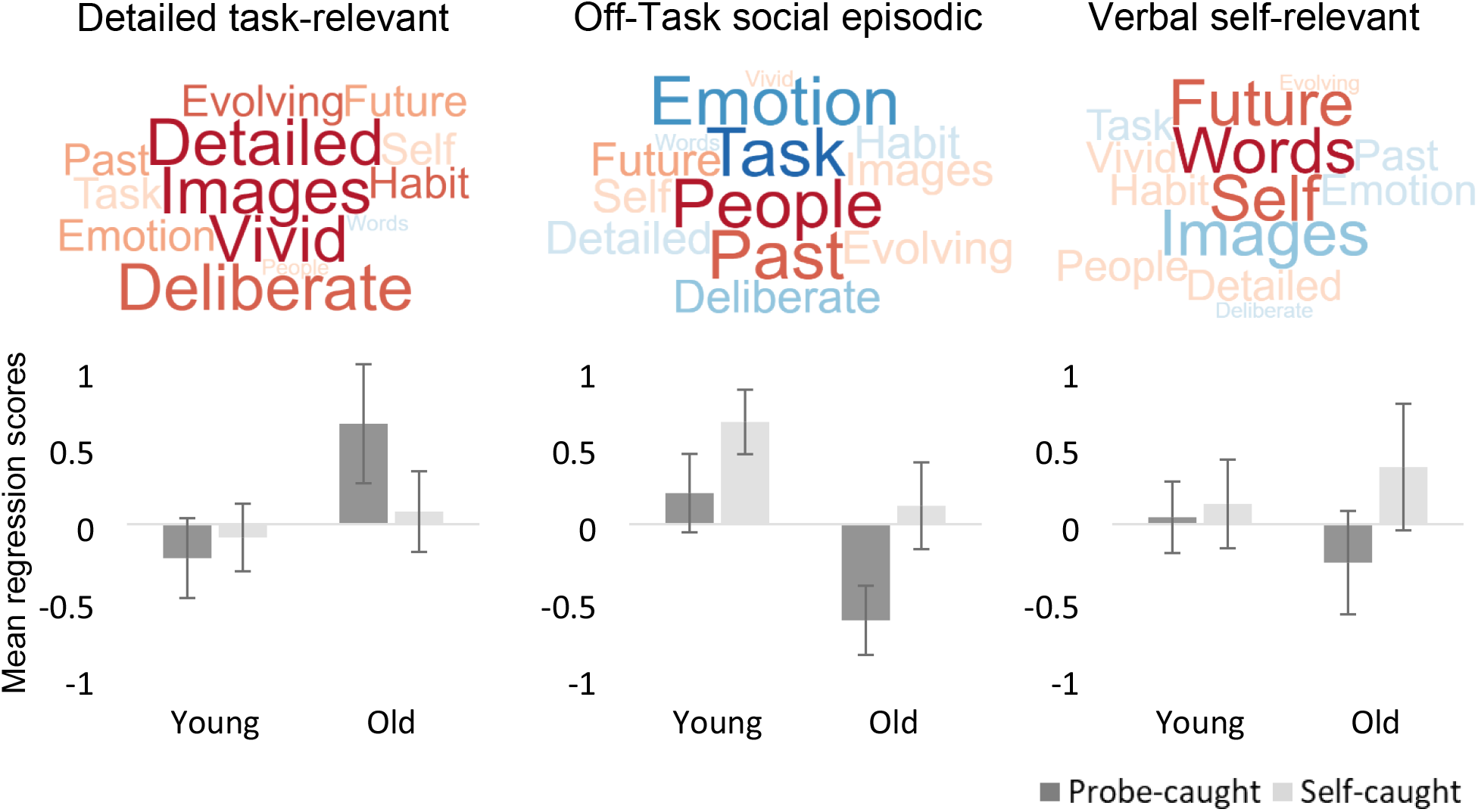
Effects of Age (*Young, Old*) and Task measures (*Probe-caught, Self-caught*) on mean regression scores of self-generated thoughts. Word clouds illustrate the loading of the different questions on the three factors resulting from the principal component analyses (PCA). Varimax rotation was applied to the data set. Error bars represent 98.3% Confidence Intervals Bonferroni corrected for the number of factors.

Analysis of **Detailed task-related thoughts** demonstrated a main effect of the type of measure with anecdotal Bayesian evidence in favour of this effect, indicating that relative to the self-caught method (*M*=-.01, *SD*=.53), experiences reported at probes captured more detailed on-task patterns (*M*=.17, *SD*=.82), *F*(1,61)=10.16, *p*=.002, *η*_*p*_ ^2^=.14, BF_10_ =1.602, BF_01_ =.624. Furthermore, we found a main effect of age with robust Bayesian evidence in favour of this effect, indicating that experiences were more detailed for older adults (*M*=.23, *SD*=.84) than younger adults (*M*=-.19, *SD*=.56), *F*(1,61)=13.41, *p*<.001, *η*_*p*_^2^=.18, BF_10_=54.698. Additionally, an interaction effect with extreme Bayesian evidence in favour of this effect, illustrated in Figure 3, was found between age and the type of measure [*F*(1,61)=26.41, *p*<.001, *η*_*p*_^2^=.30, BF_10_=3787.36].

This indicated that patterns of detailed thoughts were more present in the probe-caught reports of older adults (*M*=.33, *SD*=.93) than of young adults (*M*=-.24, *SD*=.62) [*F*(1,72)=9.73, *p*=.003, *η*_*p*_ ^2^=.12, BF_10_ =13.672], but not when measured by the self-caught method [*F*(1,61)=1.57, *p*=.215, *η*_*p*_ ^2^=.03, BF_10_=0.500]. Furthermore, thoughts had more detailed features when older individuals reported experiences at probes (*M*=.65, *SD*=.81) than when their experiences were self-caught (*M*=.08, *SD*=.54), *F*(1, 27)=21.96, *p*<.001, *η*_*p*_ ^2^=.45, BF_10_ =329.78. See Table S5 in the supplementary materials for a comparison of the models in a Bayesian repeated-measure ANOVA. These data suggest that experiential reports at probes contain more evidence of patterns of highly detailed task processing when described at probes, especially for older individuals.

Analysis of patterns of **Off-task social episodic thoughts** revealed a main effect of measure with extreme Bayesian evidence in favour of this effect, showing that on-task thought patterns were more strongly captured by the probe-caught method (*M*=-.17, *SD*=.68) while experiences had more social episodic features when they were self-caught (*M*=.42, *SD*=.60), *F*(1,61)=68.22, *p*<.001, *η*_*p*_ ^2^=.53, BF_10_ =1.416e+8. In addition, we found a main effect of age with extreme Bayesian evidence in favour of this effect, showing that patterns of on-task thought were more present in the older adults’ reports (*M*=-.53, *SD*=.47). In contrast, off-task reports were more prevalent in the reports of young adults (*M*=.35, *SD*=.53), *F*(1,61)=34.81, *p*<.001, *η*_*p*_^2^=.36, BF_10_=29889.12. However, the interaction effect between age and the type of measure was not significant when controlling for the number of comparisons with anecdotal Bayesian evidence in favour of this effect [*F*(1,61)=3.73, *p*=.058, *η*_*p*_^2^=.06; BF_10_=1.20, BF_01_=0.833]. See Table S6 in the supplementary materials for a comparison of the models in a Bayesian repeated-measure ANOVA.

Results for **Off-task verbal self-relevant thoughts** revealed a main effect of measure with strong Bayesian evidence in favour of this effect, suggesting that experiences reported at probes had fewer off-task verbal features (*M*=-.09, *SD*=.63) than experiences that were self-caught (*M*=.24, *SD*=.77), *F*(1,61)=15.44, *p*<.001, *η*_*p*_^2^=.20, BF_10_ =26.810. There was no main effect of age on this type of thought, *F*(1,61)=.033, *p*=.856, *η*_*p*_ ^2^=.00, BF_10_ =0.268. However, we found evidence of an interaction effect (See Figure 3) between age and the type of measure with anecdotal Bayesian evidence in favour of this effect, *F*(1,61)=8.80, *p*=.004, *η* _*p*_^2^=.13, BF_10_ =2.53, BF_01_=0.115). Further analyses showed that, in the case of older adults, verbal self-relevant thought patterns were less present in the probe-caught method (*M*=-.25, *SD*=.70) than when experiences were reported via the self-caught method (*M*=.37, *SD*=.86), *F*(1,27)=17.16, *p*<.001, *η*_*p*_ ^2^=.39, BF_10_ =106.13. However, this difference was not evident in the group of young adults, *F*(1,34)=.650, *p*=.426, *η*_*p*_^2^=.02, BF_10_ =0.321. See Table S7 in the supplementary materials for a comparison of the models in a Bayesian repeated-measure ANOVA.

### Interim summary

Collectively analyzing the omnibus data supports the hypothesized utility of self-caught experiences as providing insight into transient off-task states. As indicated by PCA, the dimensional structure of our experiential data highlights two dimensions of off-task thought and a single dimension of detailed task focus. We found that self-caught experiences generally captured more features related to off-task states than was seen at probes and that patterns of detailed task focus are most likely captured at probes, especially for older individuals. This pattern is consistent with the assumption that when experiences are reported because people notice a change in their experience, these reports provide insight into features of experience linked to off-task states relative to the patterns seen in self-reports at externally prompted probes.

### Relationship between thought patterns, method of report, and motivation

Our final analysis examined whether the features of experiences and their relationship to the different methods of experience sampling are related to motivation. We first examined self-reported motivation across the different cells of our analysis using a 2 (Age; Young, Old) by 2 (Measure; Probe-caught, Self-caught) repeated-measure ANOVA. This revealed a significant main effect of age, with extreme Bayesian evidence in favour of this effect, suggesting that older adults (*M*=6.34, *SD*=.89) were more motivated to complete the task than young adults (*M*=5.18, *SD*=1.06), *F*(1,72)=26.17, *p*<.001, *η*_*p*_ ^2^=.27; BF_10_ =6220.732. The effect of measures and the interaction between measures and age were not significant (*p*=.944, BF_10_=0.175 and *p*=.271, BF_10_=0.408, respectively). See Table S8 in the supplementary materials for a comparison of the models in a Bayesian repeated-measure ANOVA.

To understand how changes in thought patterns assessed by different measures and in different age groups are associated with variation in self-reported motivation levels, we ran mediation analyses following the procedure advocated by Preacher and Hayes (2004). We used the most recent version of PROCESS (version 3.5) for SPSS 26.0 (Field, 2018). We selected the following specifications: model number 4, 5000 bootstrap samples, and 95% confidence intervals. Based on our results, age-related differences were found in probe-caught detailed task-relevant thoughts, probe-caught off-task social episodic thoughts, and self-caught off-task social episodic thoughts. Therefore, each of these three outcome variables will be considered for mediation analyses, as follow (1) Age → Motivation (probe-caught) → Detailed (probe-caught), (2) Age → Motivation (probe-caught) → Social episodic (probe-caught), and (3) Age → Motivation (self-caught) → Social episodic (self-caught). These analyses used age as a continuous variable to gain specificity. For completeness, *Pearson* correlations have been conducted between the predictor (i.e., age) the outcome (i.e., detailed or social episodic thoughts), and the mediator (i.e., motivation). These are presented in Table 3.

**Table 3.**
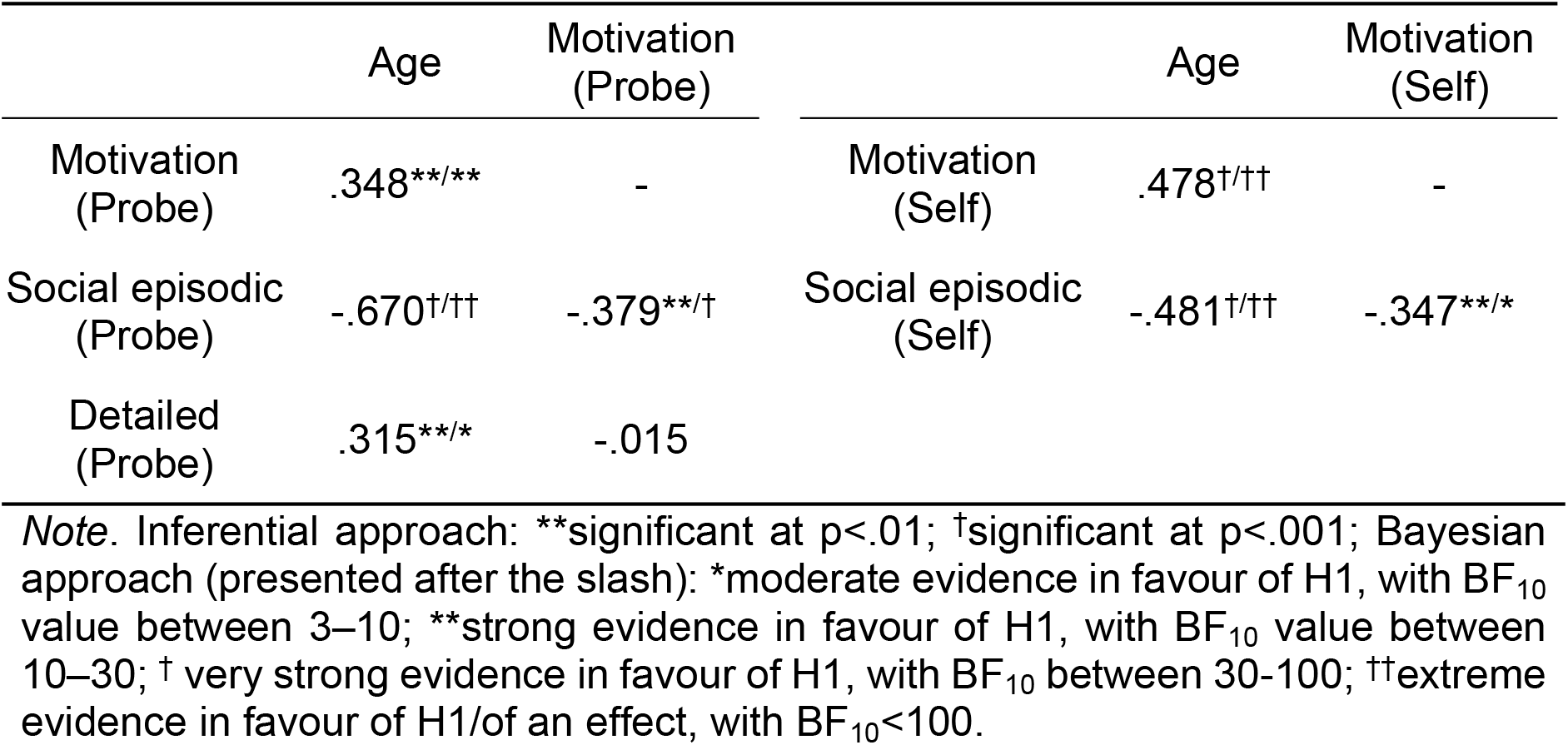
Pearson’s correlation between age, motivation rates, detailed task-relevant thoughts, and off-task social episodic thoughts collected with the probe-caught and self-caught method separately.

First, the (1) Age → Motivation (probe-caught) → Detailed (probe-caught) mediation was considered. The total effect of age on Detailed (probe) was significant, *b*=.011, CI95% [.003, .018], *p*=.006. However, the standardized indirect effect included 0 in the confidence interval, *b*=-.049, CI95% [-.155, .029], suggesting that motivation did not mediate the age-related increase of probe-caught detailed thoughts (Figure 4.). Additionally, the direct effect of age on detailed thoughts remained significant, *b*=.012, CI95% [.004, .020], *p*=.003. This analysis establishes that detailed experiences varied with age when assessed by the probe-caught method and that this difference could not be attributed to motivation differences.

**Figure 4.**
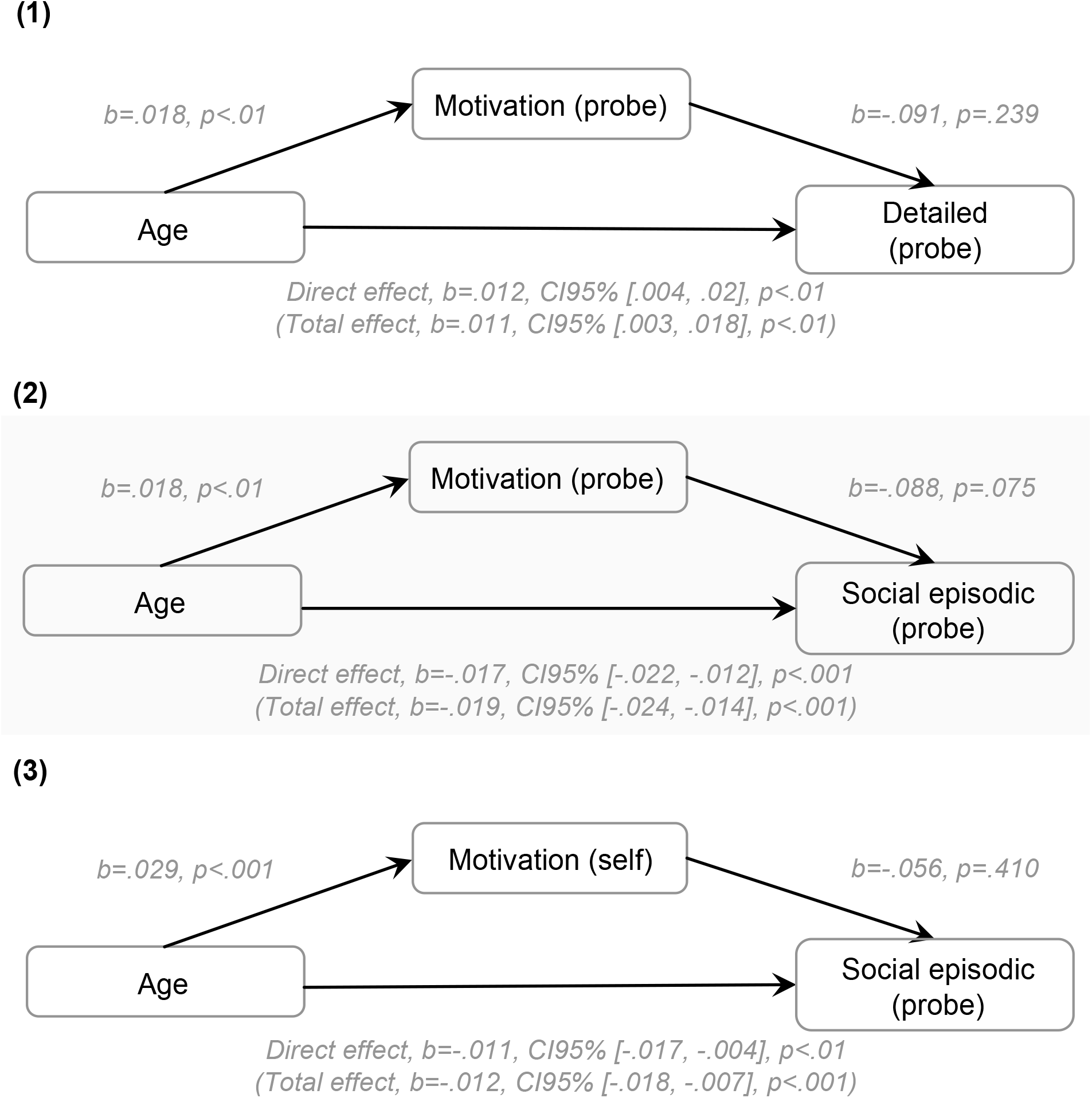
Models of age as a predictor of (1) probe-caught detailed task-relevant thoughts (2) probe-caught off-task social episodic thoughts and (3) self-caught off-task social episodic thoughts, mediated by motivation to the task.

Then, the (2) Age → Motivation (probe-caught) → Social episodic (probe-caught) mediation was conducted. The total effect of age on social episodic thoughts (probe) was significant, *b*=-.019, CI95% [-.024, -.014], *p*<.001. The standardized indirect effect did not include 0 in the confidence interval, *b*=-.058, CI95% [-.150, -.002], suggesting that an age-related increase in motivation mediated the age-related decrease in probe-caught off-task social episodic thoughts (Figure 4.). However, the direct effect of age on off-task social episodic thoughts remained significant, *b*=-.017, CI95% [-.022, -.012], *p*<.001, indicating partial mediation. This indicates that although motivation was associated with more off-task social episodic experiences when captured by probes, this difference did not fully explain age-related differences in this experience.

Finally, the (3) Age → Motivation (self-caught) → Social episodic (self-caught) mediation was tested. The total effect of age on social episodic thoughts (self) was significant, *b*=-.012, CI95% [-.018, -.007], *p*<.001. However, the standardized indirect effect included 0 in the confidence interval, *b*=-.063, CI95% [-.246, .093], suggesting that motivation did not mediate the age-related decrease of self-caught off-task social episodic thoughts (Figure 4.). Additionally, the direct effect of age on off-task social episodic thoughts remained significant, *b*=-.011, CI95% [-.017, -.004], *p*=.003. This suggests that motivation differences were neither related to age-related differences in patterns of off-task social episodic thoughts gained by the self-caught method, nor did this explain age-related differences in the patterns of thoughts.

## Discussion

Variations in ongoing thought patterns are an essential part of our everyday lives, yet, our understanding of these processes is still in its infancy, partly because of their transient and internal nature. Here, we examined the current view that features of off-task states are particularly captured by acquiring reports of experience when individuals notice changes in their ongoing conscious experiences (Schooler, 2002). Specifically, we combined two methods that are increasingly used: multi-dimensional experience sampling (Smallwood et al., 2016), in which experience is characterized along many dimensions, and through comparing experience sampling data generated by probe- and self-caught methods. We applied Principal Components Analysis to these data to provide low dimensional descriptions of their self-reports and to allow us to use a pattern learning perspective to understand the degree to which these measures tap into common underlying states.

We found three patterns of thought often found in the literature: detailed task-relevant experiences, off-task social episodic thoughts, and verbal self-relevant thoughts (Ho et al., 2019; Konu, Turnbull, et al., 2020; Sormaz et al., 2018; Turnbull et al., 2019). First, we examined the degree of similarity in the features of experiences captured using both approaches. In general, we found significant overlap between these patterns, which were clearest using the probe-caught method, and for younger relative to older individuals. The observation that probe-caught experience sampling captures ongoing thought patterns more robustly, is consistent with the notion that this experience sampling method provides a more rounded description of conscious experience because it does not depend to the same extent on self-monitoring ability (Schooler, 2002).

Importantly, our analysis also demonstrated that asking individuals to report when they noticed transient changes in experience is especially able to highlight features of experiences associated with off-task states. For example, when people noticed a change in their experience, their reports contained fewer features of detailed, task-relevant thought than when prompted by externally motivated probes. Additionally, self-caught reports contained greater evidence of both off-task thought patterns (episodic-social and verbal self-relevant) than were seen in reports collected at probes. Our pattern analysis of self-reports captured via both methods, therefore, supports the hypothesis that individuals can selectively monitor for the occurrence of specific transient changes in experience (e.g. off-task episodes).

An important concern in the present work was whether differences in transient changes in thought patterns revealed by both methods change with age. Consistent with prior studies, older adults reported fewer off-task thoughts than younger adults (Jordão et al., 2019; Maillet & Schacter, 2016). Furthermore, we confirmed prior studies with older adults reporting being more motivated to perform the task than young adults, regardless of the sampling method (Shake et al., 2016). However, our data provide constraints on why experience changes as we age. Our study found that off-task episodes can vary in modality since PCA identified two broad dimensions of off-task thoughts that were distinguished by their loading on images and words. Critically, these changed with age, with older individuals less likely to report off-task experiences, including imagery, but instead reported off-task experiences with verbal features. This is consistent with the view that visuospatial cognition is more impaired than verbal in ageing (Jenkins et al., 2000). However, data dismiss the notion that age-related changes in cognition result from deficits in meta-awareness: We found that older individuals’ self-caught episodes contained greater evidence of transient states of experience, in this case, off-task thoughts with verbal features. Second, we found that consistent with prior studies, older adults reported greater detailed task-relevant thoughts, as well as a reduction in off-task episodes (Martinon et al., 2019). Although motivation partially mediated the age-related decrease in probe-caught off-task thoughts, it did not explain age-related increases in detailed-task focus. This suggests that motivational differences can not account entirely for changes in ongoing experience with age. Critically, past research has repeatedly demonstrated increased detailed task-related experiences during more demanding tasks (Konu, Mckeown, et al., 2020; Sormaz et al., 2018; Turnbull et al., 2019). Mediation analysis identified a significant contribution of motivation in explaining the age-related gap in off-task social episodic thoughts, particularly for probe-caught episodes, a pattern consistent with a growing number of studies (Frank et al., 2015; Krawietz et al., 2012; Seli et al., 2020; Shake et al., 2016). Therefore, our data suggest that the emphasis on detailed task-focus in older adults is not likely to be related to motivational differences in our sample.

To conclude, we adopted a pattern learning approach to determine whether reports provided when participants notice changes in their experience contain features related to off-task states in a way different from those reported at probes (Schooler, 2002). We found that while self-caught episodes provide more “noisy” descriptions of experience than reports at probes, they are more likely to describe features of experience that are associated with off-task states. Nonetheless, there are important open questions that remain unanswered. First, these data do not constrain accounts of the cognitive processes that underpin the capacity for self-recognition of off-task states. Moving forward, to address this important question, it is possible that experiential features that are preferentially expressed in self-caught experiences could provide insight into if and how meta-awareness can help control conscious experience (Schooler et al., 2011). Second, while our data largely rule out the possibility that older individuals are unable to self-detect off-task thought patterns, it does not constrain accounts of why individuals tend to show less evidence of off-task thought in older age. It is worthwhile noting, however, that ten older adults (compared to 1 younger) did not self-catch thoughts, highlighting the age variability in the dataset. The extra demands during the self-catch condition mirror similar tasks employed in the prospective memory literature, where age effects are evident (Gonneaud et al. 2011). Regardless, it remains an open question the extent to which features of age-related changes in experience are primarily driven by organic changes in cognition, lifestyle differences, or familiarity with university settings (e.g. McVay et al., 2013). Increased verbal features in off-task experiences for older individuals may be related to the increased reliance on semantic knowledge and crystallized intelligence as we grow older (e.g. Riby et al. 2004), which may have links to organic changes in neural function (Spreng et al., 2018). This possibility is consistent with the abovementioned suggestion of greater declines of visuospatial ability as we age and could be explored by combining self- and probe-caught experience sampling methods in older and younger individuals in conjunction with fMRI measures (Kucyi, 2018). Additionally, thoughts’ temporal patterns may bring further perspective to understanding these age-related variabilities (Zanesco, 2020). Third, our study leveraged a pattern learning approach to determine how different methods capture different features of experience, however, this observation does not overcome the need for better ground truth in this line of research. A complete account of these states is likely to require more refined psychological theories and better indirect markers of hidden private states than are available at present. Our study is an important step toward this goal since it illustrates how similarities and differences in the patterns generated by different self-report methods can help interrogate psychological accounts of experience.

## Open Practices

Data and supplementary material can be found at https://figshare.northumbria.ac.uk/ [complete link to be inserted by corresponding author on acceptance]

## Footnotes

1 BF_10_ corresponds to the Bayesian probability of the occurrence of a hypothesis (H1) and the likelihood of another null hypothesis (H0). It was calculated with JASP 0.14.1.0 (JASP Team, 2017) and interpreted according to Lee and Wagenmakers (2014 adjusted from Jeffreys, 1961). All priors were equal.

2 As recommended, and for the rest of the manuscript, BF_10_ corresponding to the interaction and both main effects was divided by BF_10_ of both main effects.

## References

Allen, M., Smallwood, J., Christensen, J., Gramm, D., Rasmussen, B., Gaden Jensen, C., Roepstorff, A., & Lutz, A. (2013). The balanced mind : The variability of task-unrelated thoughts predicts error-monitoring. Frontiers in Human Neuroscience, 7. https://doi.org/10.3389/fnhum.2013.00743

Barron, E., Riby, L. M., Greer, J., & Smallwood, J. (2011). Absorbed in thought: The effect of mind wandering on the processing of relevant and irrelevant events. Psychological science, 22(5), 596–601.

Damoiseaux, J. S. (2017). Effects of aging on functional and structural brain connectivity. NeuroImage, 160, 32–40. https://doi.org/10.1016/j.neuroimage.2017.01.077

Ennis, G. E., Hess, T. M., & Smith, B. T. (2013). The Impact of Age and Motivation on Cognitive Effort : Implications for Cognitive Engagement in Older Adulthood. Psychology and aging, 28(2), 495–504. https://doi.org/10.1037/a0031255

Field, A. (2018). Discovering Statistics Using IBM SPSS Statistics 5th ed. Sage. http://dln.jaipuria.ac.in:8080/jspui/handle/123456789/812

Fjell, A. M., Walhovd, K. B., Fennema-Notestine, C., McEvoy, L. K., Hagler, D. J., Holland, D., Brewer, J. B., & Dale, A. M. (2009). One-Year Brain Atrophy Evident in Healthy Aging. Journal of Neuroscience, 29(48), 15223–15231. https://doi.org/10.1523/JNEUROSCI.3252-09.2009

Folstein, M. F., Folstein, S. E., & McHugh, P. R. (1975). “Mini-mental state”. Journal of Psychiatric Research, 12(3), 189–198. https://doi.org/10.1016/0022-3956(75)90026-6

Frank, D. J., Nara, B., Zavagnin, M., Touron, D. R., & Kane, M. J. (2015). Validating older adults’ reports of less mind-wandering : An examination of eye movements and dispositional influences. Psychology and Aging, 30(2), 266–278. https://doi.org/10.1037/pag0000031

Gonneaud, J., Kalpouzos, G., Bon, L., Viader, F., Eustache, F., & Desgranges, B. (2011). Distinct and shared cognitive functions mediate event-and time-based prospective memory impairment in normal ageing. Memory, 19(4), 360–377.

Ho, N. S. P., Poerio, G., Konu, D., Turnbull, A., Sormaz, M., Leech, R., Bernhardt, B., Jefferies, E., & Smallwood, J. (2020). Facing up to the wandering mind : Patterns of off-task laboratory thought are associated with stronger neural recruitment of right fusiform cortex while processing facial stimuli. NeuroImage, 214, 116765. https://doi.org/10.1016/j.neuroimage.2020.116765

Ho, N. S. P., Wang, X., Vatansever, D., Margulies, D. S., Bernhardt, B., Jefferies, E., & Smallwood, J. (2019). Individual variation in patterns of task focused, and detailed, thought are uniquely associated within the architecture of the medial temporal lobe. NeuroImage, 202, 116045. https://doi.org/10.1016/j.neuroimage.2019.116045

Jenkins, L., Myerson, J., Joerding, J. A., & Hale, S. (2000). Converging evidence that visuospatial cognition is more age-sensitive than verbal cognition. Psychology and Aging, 15(1), 157–175. https://doi.org/10.1037/0882-7974.15.1.157

Jordano, M. L., & Touron, D. R. (2017). Stereotype threat as a trigger of mind-wandering in older adults. Psychology and Aging, 32(3), 307–313. https://doi.org/10.1037/pag0000167

Jordão, M., Ferreira-Santos, F., Pinho, M. S., & St. Jacques, P. L. (2019). Meta-analysis of aging effects in mind wandering : Methodological and sociodemographic factors. Psychology and Aging. https://doi.org/10.1037/pag0000356

Karapanagiotidis, T., Bernhardt, B. C., Jefferies, E., & Smallwood, J. (2017). Tracking thoughts : Exploring the neural architecture of mental time travel during mind-wandering. NeuroImage, 147, 272–281. https://doi.org/10.1016/j.neuroimage.2016.12.031

Konishi, M., McLaren, D. G., Engen, H., & Smallwood, J. (2015). Shaped by the Past : The Default Mode Network Supports Cognition that Is Independent of Immediate Perceptual Input. PLOS ONE, 10(6), e0132209. https://doi.org/10.1371/journal.pone.0132209

Konu, D., Mckeown, B., Turnbull, A., Ho, N. S. P., Vanderwal, T., McCall, C., Tipper, S. P., Jefferies, E., & Smallwood, J. (2020). Exploring patterns of ongoing thought under naturalistic and task-based conditions. BioRxiv, 2020.07.29.226431. https://doi.org/10.1101/2020.07.29.226431

Konu, D., Turnbull, A., Karapanagiotidis, T., Wang, H.-T., Brown, L. R., Jefferies, E., & Smallwood, J. (2020). A role for the ventromedial prefrontal cortex in self-generated episodic social cognition. NeuroImage, 218, 116977. https://doi.org/10.1016/j.neuroimage.2020.116977

Krawietz, S. A., Tamplin, A. K., & Radvansky, G. A. (2012). Aging and mind wandering during text comprehension. Psychology and Aging, 27(4), 951–958. https://doi.org/10.1037/a0028831

Kucyi, A. (2018). Just a thought : How mind-wandering is represented in dynamic brain connectivity. NeuroImage, 180, 505–514. https://doi.org/10.1016/j.neuroimage.2017.07.001

Lee, M. D., & Wagenmakers, E.-J. (2014). Bayesian Cognitive Modeling : A Practical Course. Cambridge University Press.

Maillet, D., & Schacter, D. L. (2016). From mind wandering to involuntary retrieval : Age-related differences in spontaneous cognitive processes. Neuropsychologia, 80, 142–156. https://doi.org/10.1016/j.neuropsychologia.2015.11.017

Martinon, L. M., Riby, L. M., Poerio, G., Wang, H.-T., Jefferies, E., & Smallwood, J. (2019). Patterns of on-task thought in older age are associated with changes in functional connectivity between temporal and prefrontal regions. Brain and Cognition, 132, 118–128. https://doi.org/10.1016/j.bandc.2019.04.002

McVay, J. C., & Kane, M. J. (2010). Does mind wandering reflect executive function or executive failure? Comment on Smallwood and Schooler (2006) and Watkins (2008). Psychological Bulletin, 136(2), 188–197. https://doi.org/10.1037/a0018298

McVay, J. C., Meier, M. E., Touron, D. R., & Kane, M. J. (2013). Aging ebbs the flow of thought : Adult age differences in mind wandering, executive control, and self-evaluation. Acta Psychologica, 142(1), 136–147. https://doi.org/10.1016/j.actpsy.2012.11.006

Peirce, J. W. (2007). PsychoPy—Psychophysics software in Python. Journal of Neuroscience Methods, 162(1), 8–13. https://doi.org/10.1016/j.jneumeth.2006.11.017

Peugh, J. L., & Enders, C. K. (2005). Using the SPSS Mixed Procedure to Fit Cross-Sectional and Longitudinal Multilevel Models. Educational and Psychological Measurement, 65(5), 717–741. https://doi.org/10.1177/0013164405278558

Preacher, K. J., & Hayes, A. F. (2004). SPSS and SAS procedures for estimating indirect effects in simple mediation models. Behavior Research Methods, Instruments, & Computers, 36(4), 717–731. https://doi.org/10.3758/BF03206553

Riby, L. M., Perfect, T. J., & Stollery, B. T. (2004). Evidence for disproportionate dual-task costs in older adults for episodic but not semantic memory. Quarterly Journal of Experimental Psychology Section A, 57(2), 241–267.

Sayette, M. A., Reichle, E. D., & Schooler, J. W. (2009). Lost in the Sauce : The Effects of Alcohol on Mind Wandering. Psychological Science, 20(6), 747–752. https://doi.org/10.1111/j.1467-9280.2009.02351.x

Sayette, M. A., Schooler, J. W., & Reichle, E. D. (2010). Out for a Smoke : The Impact of Cigarette Craving on Zoning Out During Reading. Psychological Science, 21(1), 26–30. https://doi.org/10.1177/0956797609354059

Schooler, J. W. (2002). Re-representing consciousness : Dissociations between experience and meta-consciousness. Trends in Cognitive Sciences, 6(8), 339–344. https://doi.org/10.1016/S1364-6613(02)01949-6

Schooler, J. W., Smallwood, J., Christoff, K., Handy, T. C., Reichle, E. D., & Sayette, M. A. (2011). Meta-awareness, perceptual decoupling and the wandering mind. Trends in Cognitive Sciences. https://doi.org/10.1016/j.tics.2011.05.006

Seli, P., O’Neill, K., Carriere, J. S. A., Smilek, D., Beaty, R. E., & Schacter, D. L. (2020). Mind-Wandering Across the Age Gap : Age-Related Differences in Mind-Wandering Are Partially Attributable to Age-Related Differences in Motivation. The Journals of Gerontology: Series B, gbaa031. https://doi.org/10.1093/geronb/gbaa031

Shake, M. C., Shulley, L. J., & Soto-Freita, A. M. (2016). Effects of Individual Differences and Situational Features on Age Differences in Mindless Reading. The Journals of Gerontology Series B: Psychological Sciences and Social Sciences, 71(5), 808–820. https://doi.org/10.1093/geronb/gbv012

Smallwood, J., Karapanagiotidis, T., Ruby, F., Medea, B., Caso I. de, Konishi, M., Wang, H.-T., Hallam, G., Margulies, D. S., & Jefferies, E. (2016). Representing Representation : Integration between the Temporal Lobe and the Posterior Cingulate Influences the Content and Form of Spontaneous Thought. PLOS ONE, 11(4), e0152272. https://doi.org/10.1371/journal.pone.0152272

Smallwood, J., & Schooler, J. W. (2006). The restless mind. Psychological Bulletin, 132(6), 946–958. https://doi.org/10.1037/0033-2909.132.6.946

Smallwood, J., Turnbull, A., Wang, H., Ho, N. S. P., Poerio, G. L., Karapanagiotidis, T., Konu, D., Mckeown, B., Zhang, M., Murphy, C., Vatansever, D., Bzdok, D., Konishi, M., Leech, R., Seli, P., Schooler, J. W., Bernhardt, B., Margulies, D. S., & Jefferies, E. (2021). The neural correlates of ongoing conscious thought. IScience, 24(3), 102132. https://doi.org/10.1016/j.isci.2021.102132

Sormaz, M., Murphy, C., Wang, H., Hymers, M., Karapanagiotidis, T., Poerio, G., Margulies, D. S., Jefferies, E., & Smallwood, J. (2018). Default mode network can support the level of detail in experience during active task states. Proceedings of the National Academy of Sciences, 115(37), 9318–9323. https://doi.org/10.1073/pnas.1721259115

Spreng, R. N., Lockrow, A. W., DuPre, E., Setton, R., Spreng, K. A. P., & Turner, G. R. (2018). Semanticized autobiographical memory and the default–executive coupling hypothesis of aging. Neuropsychologia, 110, 37–43. https://doi.org/10.1016/j.neuropsychologia.2017.06.009

Turnbull, A., Wang, H. T., Murphy, C., Ho, N. S. P., Wang, X., Sormaz, M., Karapanagiotidis, T., Leech, R. M., Bernhardt, B., Margulies, D. S., Vatansever, D., Jefferies, E., & Smallwood, J. (2019). Left dorsolateral prefrontal cortex supports context-dependent zprioritization of off-task thought. Nature Communications, 10(1), 3816. https://doi.org/10.1038/s41467-019-11764-y

West, R. L. (2000). In defense of the frontal lobe hypothesis of cognitive aging. Journal of the International Neuropsychological Society, 6(6), 727–729.

Zanesco, A. P. (2020). Quantifying streams of thought during cognitive task performance using sequence analysis. Behavior Research Methods, 52(6), 2417–2437. https://doi.org/10.3758/s13428-020-01416-1

